# Spinal interneuron population dynamics underlying flexible pattern generation

**DOI:** 10.1101/2024.06.20.599927

**Authors:** Lahiru N. Wimalasena, Chethan Pandarinath, Nicholas Au Yong

**Affiliations:** Wallace H. Coulter Department of Biomedical Engineering, Emory University and Georgia Institute of Technology, Atlanta, GA, USA; Department of Neurosurgery, Emory University, Atlanta, GA, USA; Department of Cell Biology, Emory University, Atlanta, GA, USA

## Abstract

The mammalian spinal locomotor network is composed of diverse populations of interneurons that collectively orchestrate and execute a range of locomotor behaviors. Despite the identification of many classes of spinal interneurons constituting the locomotor network, it remains unclear how the network’s collective activity computes and modifies locomotor output on a step-by-step basis. To investigate this, we analyzed lumbar interneuron population recordings and multi-muscle electromyography from spinalized cats performing air stepping and used artificial intelligence methods to uncover state space trajectories of spinal interneuron population activity on single step cycles and at millisecond timescales. Our analyses of interneuron population trajectories revealed that traversal of specific state space regions held millisecond-timescale correspondence to the timing adjustments of extensor-flexor alternation. Similarly, we found that small variations in the path of state space trajectories were tightly linked to single-step, microvolt-scale adjustments in the magnitude of muscle output.

**One sentence summary:** Features of spinal interneuron state space trajectories capture variations in the timing and magnitude of muscle activations across individual step cycles, with precision on the scales of milliseconds and microvolts respectively.

## Introduction

The nervous system can generate a wide repertoire of movements through precisely coordinating the time-varying activity of muscles. Perhaps the most well-studied example of this is locomotion, where central pattern generating (CPG) circuits intrinsic to the lumbar spinal cord activate flexor and extensor muscles in an alternating pattern to execute full step cycles (Frigon, 2012; Grillner and El Manira, 2020). Flexibility in coordinating when, and how much, to activate flexor and extensor muscles is crucial for maintaining stable locomotion, ensuring that the foot is appropriately lifted, securely planted, and able to drive the body forward with each step. The spinal cord’s capacity to autonomously produce and adjust these muscle activation patterns has been known for over a century (Brown, 1911; Sherrington, 1906). Many anatomically- and genetically-identified classes of spinal interneurons play a critical role in mammalian locomotion (Goulding, 2009); however, it is still unclear how these different classes of interneurons, acting collectively as a network, perform the appropriate computations necessary to produce the alternating patterns of muscle activations that underpin different locomotor behaviors.

A popular approach to understand the network-level function of mammalian spinal interneurons has focused on simulating the activity of posited networks of interneurons involved in the locomotor circuit (McCrea and Rybak, 2008; Rybak et al., 2015). Groups of distinct interneuron populations are wired together in an experimenter-defined connectivity pattern in order to recapitulate observed characteristics of recorded muscle activity (e.g., timing of flexor-extensor alternation) during locomotor behaviors. Such approaches put forward credible mechanistic implementations of the underlying biological circuit, which have been refined with our growing knowledge of the anatomical connections and genetic diversity that underlie the expansive spinal interneuron networks. However, this is a complex and under-constrained modeling challenge, as numerous potential connectivity patterns among interneuron populations could yield identical outputs, making the validation of such models particularly difficult. Further, continuing to adjust these simulated spinal networks to mirror our evolving understanding of interneurons is becoming progressively more difficult as the number of identified classes continues to grow (Bagnall, 2022).

An alternative perspective on studying network-level function of neural circuits can be drawn from recent approaches developed in the past two decades to interpret simultaneously-recorded neural population activity. The core idea of these approaches is that network computations can be understood by studying the distributed, dynamic patterns that have previously been shown to underlie the activity of recorded neural populations – including recent demonstrations in spinal interneurons of turtles during rhythmic scratching (Lindén et al., 2022). These distributed patterns across the population are consistent with an increasingly popular *state space* perspective on neural population activity, which posits that a coordinated dynamical system underlies the activity of individual neurons (Vyas et al., 2020). In this view, the current state of the network and its temporal evolution are the key variables that determine the circuit’s function. Viewing neural population activity through the lens of network states and their dynamics has provided new insights into neural computations underlying motor output, sensory processing, timing, decision-making, learning, and working memory.

The state space perspective provides an alternate level of abstraction from which to understand and interrogate network computations in the mammalian spinal circuit. However, it remains to be determined the degree to which spinal network states capture the temporal precision and fidelity with which the circuit shapes muscle activity, and further, which aspects of the network’s state correspond to specific features in muscle activity. To determine whether a close correspondence between interneuron population state space trajectories and generated muscle output exists, we analyzed lumbar population activity from multiple single-unit recordings along with simultaneous multi-muscle intramuscular electromyography (EMG) from two spinalized cats performing air stepping (McMahon et al., 2022).

We used artificial intelligence methods to uncover state space trajectories of the recorded interneuron population activity and EMG with high precision on the time scale of milliseconds (Keshtkaran et al., 2022; Wimalasena et al., 2022). Simultaneously recording the neuronal populations composing the locomotor circuit is technically challenging due to the anatomical spread of the involved spinal interneurons across several lumbar segments (Cazalets and Bertrand, 2000; Cowley and Schmidt, 1997). Thus to create a unified view of interneuron population activity from multiple lumbar segments, we aligned separate recordings from individual animals using dynamic neural stitching (Pandarinath et al., 2018). Analysis of the uncovered state space trajectories revealed precise links between the dominant patterns of spinal interneuron population activity and muscle activity. Specifically, we observed a regional organization in interneuron state space, i.e., traversal of particular state space locations was linked to transitions between flexor and extensor activation, with precision on the order of milliseconds. Consistent with this regional organization, we found that occasional aberrant locomotor output patterns – a phenomenon known as deletions – corresponded to a failure of the interneuron population to enter the appropriate regions of state space. And finally, we found a precise correspondence between interneuron state space trajectories and single-cycle variations in the magnitude of muscle activations.

Together, these results demonstrate that data-driven state space approaches enable the study of network interactions that underlie the nervous system’s ability to flexibly control behavior with remarkable precision. This ability to precisely interrogate network-level function may provide a useful reference when tying details of biological circuitry within the spinal cord to the computations that the circuit implements.

## Results

We analyzed population activity from multiple, sorted lumbar spinal interneurons and simultaneously recorded muscle activity from two sub-acute spinalized cats (i.e., cats were spinalized at spinal level T11/T12, 22-23 days prior to acute experiment) in an air stepping paradigm (**Fig. 1a**). To incorporate interneurons spanning the lumbar spinal cord, we combined population data from multiple air stepping sessions that sampled interneuron population activity at different locations between L3 and L7 (Cat 1: 382 neurons, Cat 2: 341 neurons).

**Figure 1.**
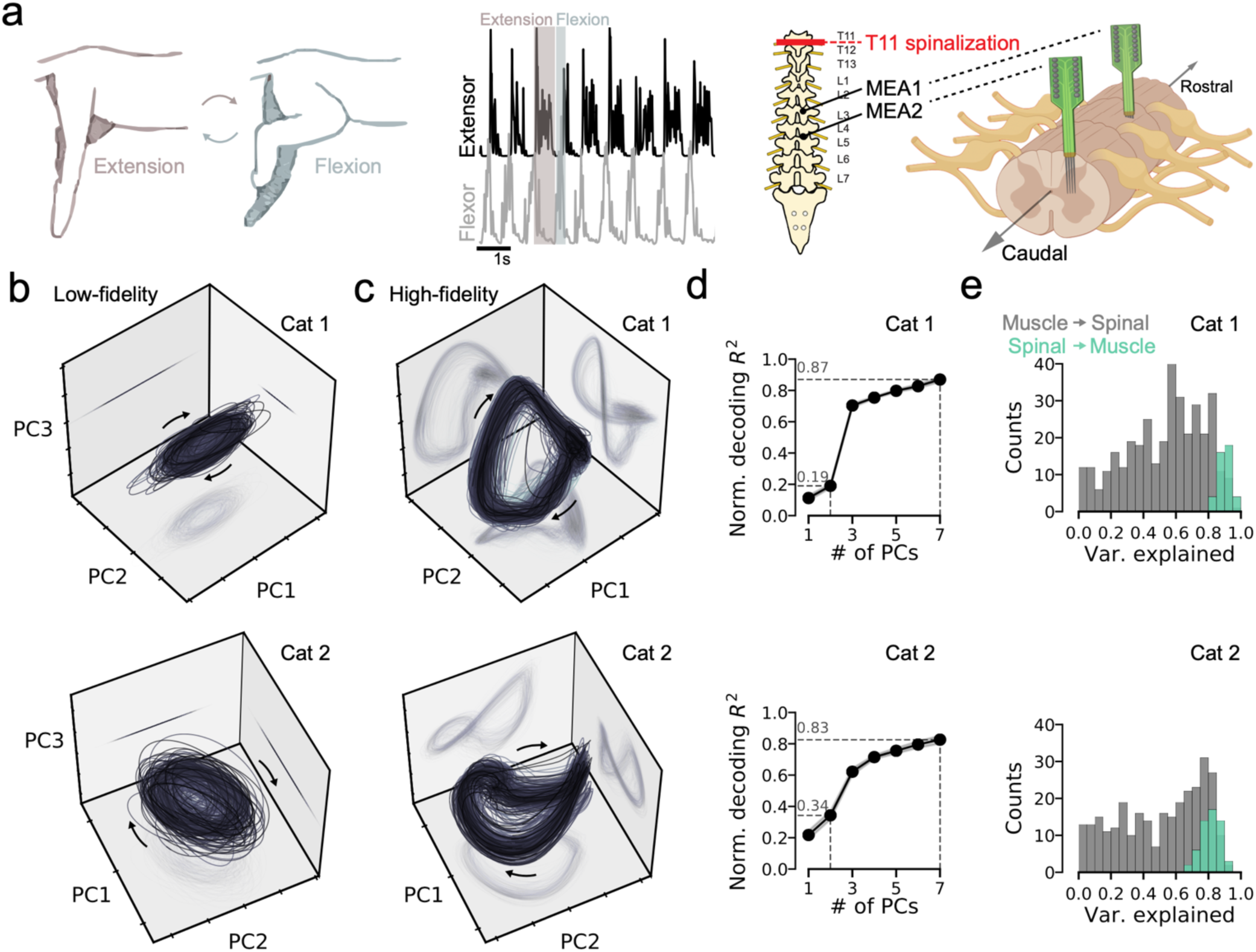
High-fidelity characterization of spinal interneuron population activity reveals state space structure beyond planar rotations. (a) Schematic of experimental setup. Sub-acute spinalized cats were administered clonidine to induce air-stepping, a rhythmic locomotor flexion and extension of the hindlimbs. Interneuron activity was recorded via two microelectrode arrays (2x64-channel probes) implanted in the intermediate zone of the right spinal hemicord. Simultaneous intramuscular EMG recordings captured activation of muscles spanning both hindlimbs (4 extensors, 3 flexors). De-noised muscle activation estimates from AutoLFADS shown for a single ipsilateral extensor muscle (biceps femoris anterior) and flexor muscle (biceps femoris posterior). (b) Single-cycle state space trajectories for spinal interneuron activity with firing rates computed through temporal smoothing (cat 1: 360 cycles, cat 2: 223 cycles). The top three principal components (PCs) are shown. (c) Same as (b), but for output factors inferred by AutoLFADS. (d) Decoding accuracy as a function of the number of spinal PCs used for muscle activation prediction. *R^2^* values are normalized to maximum prediction accuracy. (e) Histograms quantifying the proportion of explained variance for spinal interneurons’ activity predicted from muscle activations (gray) and muscle activations predicted from spinal activity (green).

We first assessed whether our data reflected similar population-level structure previously identified in interneuron population recordings from isolated turtle lumbar spinal cords during rhythmic hindlimb scratching movements (Lindén et al., 2022). We started by analyzing the tuning characteristics of the interneuron population with respect to the phase of the step cycle (**Supp. Fig. 1**). We found that the distribution of peak firing rates of individual interneurons spanned all phases of the step cycle, but skews toward the beginning and end of the step cycle (similar to phase distributions observed in some isolated turtle preparations). Given the similarities in tuning characteristics, we next asked whether our data also reflected similar state space structure (i.e., planar rotations) found in the previous study. Considering the differences in model systems (turtle vs. cat) and behaviors (scratching vs. air stepping), we wondered whether a similar rotational state space structure exists within the population activity of the mammalian locomotor network. We followed a similar protocol as (Lindén et al., 2022) and applied temporal smoothing as a means to de-noise the individual neurons’ responses and used principal components analysis (PCA) to isolate the dominant patterns distributed across the population’s activity.

We found that applying similar amounts of temporal smoothing enabled recovery of planar rotational structure (**Fig. 1b)**, however when smoothing was reduced, there was an improved moment-by-moment correspondence between smoothed interneuron population activity and muscle activity (**Supp. Fig. 2**). This highlights a limitation of using temporal smoothing to de-noise recorded neural activity: choosing the appropriate amount of smoothing involves setting an arbitrary, static cutoff on the temporal precision of neural activity features that are considered signal versus noise.

To estimate spinal interneuron firing rates more precisely, we applied AutoLFADS (Keshtkaran et al., 2022; Wimalasena et al., 2022), an unsupervised deep learning-based modeling approach. AutoLFADS infers firing rates from observed neural activity by modeling the temporal patterns that underlie the coordinated firing across neurons (i.e., dynamics) using recurrent neural networks. We trained a shared AutoLFADS model across the multiple recordings of interneuron population activity (Cat 1: 6 sessions, Cat 2: 8 sessions) from different rostro-caudal locations in the lumbar spinal cord using *dynamic neural stitching* (Pandarinath et al., 2018). Applying PCA to the AutoLFADS output revealed state-space trajectories that were more complex than the previously-reported planar rotations (**Fig. 1c**). Given the stark differences between the low-dimensional trajectories produced by the two approaches, we tested whether the firing rate estimates inferred by AutoLFADS were indeed higher-fidelity than those inferred by temporal smoothing by comparing how much information each representation contained about muscle activations. As expected, AutoLFADS-inferred firing rates enabled prediction of muscle activations with remarkable accuracy (*R^2^*>0.83 across all muscles for two cats; **Supp. Fig. 2**).

To test whether state space features inferred by AutoLFADS beyond the planar rotations were important for the prediction of muscle activity, we measured how the neural activity patterns that corresponded to muscle activations were distributed across the dominant components of the population state space trajectories. Specifically, we predicted muscle activations from spinal activity using linear regression from increasing numbers of principal components (**Fig. 1d**). Indeed, the top two principal components alone had only 20-35% of the predictive power of the full population. Adding the third PC led to a step-like increase in prediction accuracy, followed by smaller, successive gains in prediction accuracy with the inclusion of additional PCs.

Given the close anatomical proximity of spinal interneurons to motorneurons, it is conceivable that spinal interneuron population activity would be dominated by muscle-like signals. However, when we attempted to predict interneurons’ firing rates from muscle activations, there was a wide range in prediction quality: some neurons could be well-explained by muscle activations, while others were dominated by non-muscle-like patterns (**Fig. 1e**). Additionally, we found that spinal population activity was higher dimensional than muscle activity (**Supp. Fig 3**) indicating that non-muscle-like patterns contribute to the dominant components underlying the neural population. These findings are consistent with recent studies demonstrating that motor pattern-generating neural circuits contain components of population activity that do not directly reflect motor output, but are critical to supporting the pattern-generating process (i.e., *output-null dimensions*, (Churchland and Shenoy, 2024; Kaufman et al., 2014; Russo et al., 2018)).

Previous studies in motor cortical circuits have posited a variety of features of neural population state space trajectories that can be linked to muscle activity (Kaufman et al., 2014; Russo et al., 2018). However, the precision of this link has been unclear, as motor cortical circuits are anatomically far removed from spinal motorneurons, the true output of the motor system. Additionally, studies in motor cortex often use trial-averaging and temporal-smoothing of neural activity, which can limit how precisely population activity can be related to motor output. In contrast, we analyzed neural populations (spinal interneurons) that are situated in close proximity to motorneurons that drive muscle activations. Given the ability to predict muscle activity from interneuron firing rates, we aimed to determine the precision with which features of interneuron state space trajectories correlated with variations in the timing and magnitude of muscle activity. If a precise relationship existed, then this would imply that state space trajectories not only capture the circuit’s operations, but can do so at the fidelity that it produces observed behavioral variations.

We first needed to identify key ways in which the muscle activity (i.e., the circuit’s output) varied from cycle-to-cycle. Spinalized cats are known to adjust their hindlimb locomotor pattern during treadmill locomotion at varying speeds, primarily through adjusting the duration of the stance (extension) phase to produce faster steps with increases in treadmill speed (Gossard et al., 2011). We noted similar timing variability during the extension phase of the step cycle during air-stepping, where the extensor muscle’s burst duration – i.e., the interval when it was active – varied over hundreds of milliseconds depending on the length of the step cycle (**Fig. 2a**). We visualized the extensor’s muscle activity when averaged across cycles subgrouped by different durations (**Fig. 2b**) and compared it with cycle-averaged interneuron firing rates (**Fig. 2c**). As expected from our previous analyses (**Fig. 1e**), individual interneurons’ firing patterns corresponded to extensor muscle activation with varying degrees: some showed rough similarity, while others did not resemble muscle activation at all.

**Figure 2.**
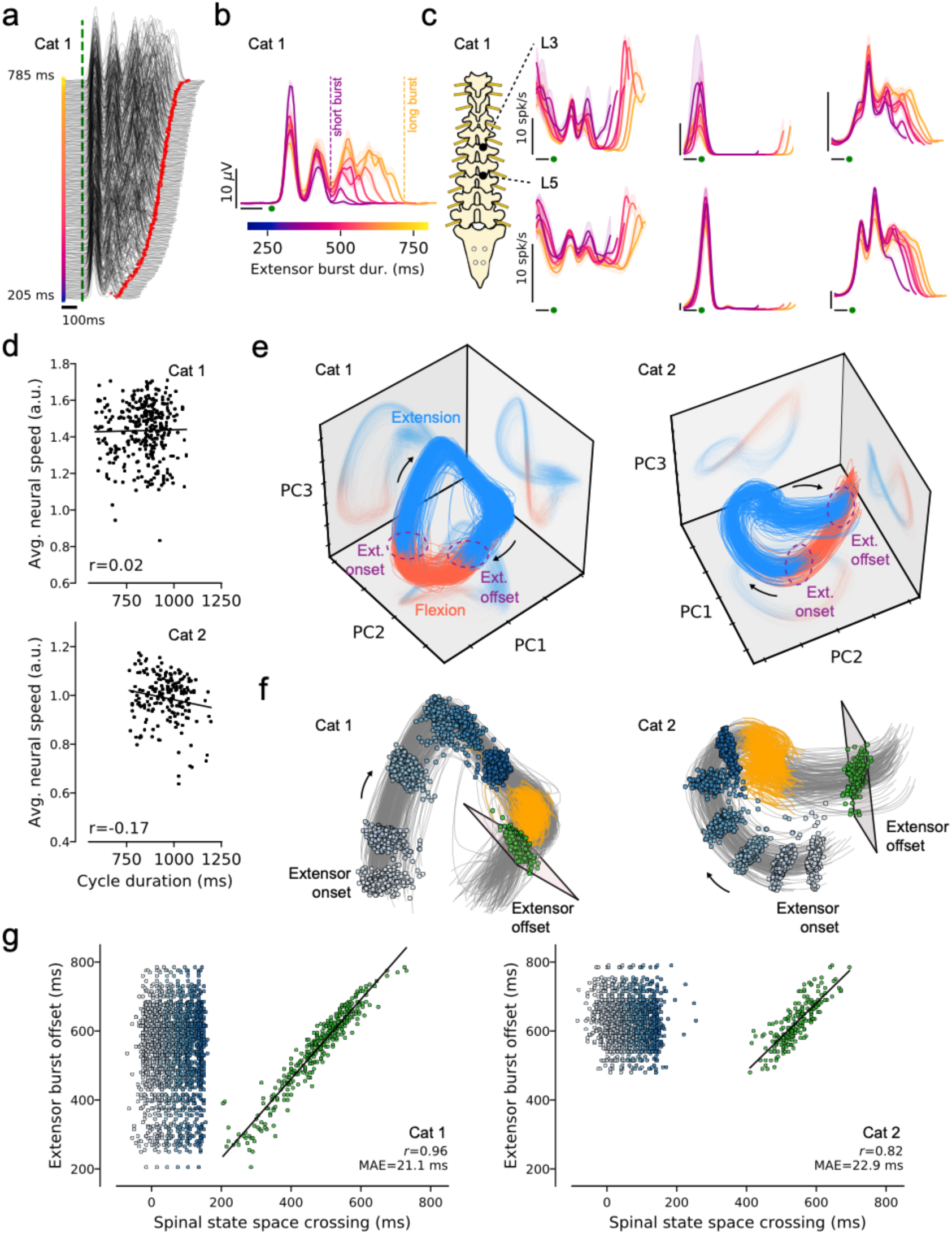
Transitions in muscle output correspond to regions in interneuron state space. (a) De-noised single-cycle muscle activations for a single ipsilateral extensor muscle (biceps femoris anterior), sorted by burst duration during extension phase of locomotion. Durations varied from 205 to 785 ms. The pulsatile nature of the extensor burst is a known EMG characteristic during clonidine-induced locomotion in spinal cats (Giroux et al., 2001; Langlet et al., 2005). (b) Extensor muscle activity averaged across cycles that were subgrouped by burst duration. Colorbar indicates the duration of the extension phase. (c) Cycle-averaged firing rates for interneurons recorded in different locations of the lumbar spinal cord. Averaging used the same cycle groupings as in (b). (d) Scatter plot comparing the average speed of state space trajectories (computed in the first 180 ms relative to extensor onset) and the duration of the gait cycle. We observed no clear linear relationship (cat 1: *r*=0.02, cat 2: *r*=-0.17). (e) Interneuron state space trajectories (top three PCs) colored based on locomotor phase (blue: extension, red: flexion). Locations in state space that correspond with transitions in extensor muscle activation marked with purple dashed circles (f) Crossing locations for 7 tangent planes along the extension region, with 6 planes positioned prior to the hold region (orange) and one plane after (seven planes were used, one example shown). Graded blue colors show crossing events for planes prior to the hold region (white: near extensor onset, dark blue: ∼160 ms after extensor onset). Green dots show crossing events for a plane after the hold region. (g) Scatter plot visualizing the relationship between the timing of state space crossing events, and the variations in extensor burst duration across individual cycles. Following the hold region, plane crossing times were strongly correlated with extensor burst offset on individual cycles (cat 1: *r*=0.96, median absolute error (MAE)=21.1 ms; cat 2: *r*=0.82, MAE=22.9 ms).

Previous work has suggested that circular or elliptical trajectories underlying neural population activity could set a basic rhythm to support pattern generation (Kirk et al., 2023; Lindén et al., 2022; Russo et al., 2018; Saxena et al., 2022). One hypothesis is that neural populations control timing by adjusting the speed of these rotational dynamics. To test whether a proportional relationship between neural population activity’s rotational speed and locomotor speed exists, we analyzed activity during extensor activation and computed neural speed as the Euclidean distance traveled by the state space trajectory between successive time steps. Perhaps surprisingly, we found little to no correlation between neural speed and cycle period for individual step cycles (**Fig. 2d**; *r*=0.02 and -0.17 for cat 1 and cat 2, respectively).

Instead, our first indication of a precise relationship between state space trajectories and the timing of muscle output came when we visualized the trajectories and simply colored them by the flexion and extension phases of the step cycle (**Fig. 2e**; phases defined by transitions in muscle activity, see **Supp. Fig. 4**). We observed a clear regional organization in state space: despite the wide variation in durations across hundreds of cycles, key muscle output transitions (i.e., flexion onset and extension onset) were consistently localized to particular regions of state space, independent of the duration of any particular cycle.

To better understand this relationship, we subgrouped cycles based on extensor burst duration and visualized the time-varying state space trajectories and corresponding flexor/extensor muscle activations for short and long duration groups (**<u>Supplementary Video 1</u>**). Trajectories dwelled at a *hold region* in state space for a duration that corresponded with variations in muscle output timing. At extension initiation, all trajectories followed a consistent arc that dominated the extension region of state space. During the extensor hold phase, state space trajectories remained in the hold region at the end of arc for a variable amount of time that appeared to correspond to the duration of extension for each cycle.

To quantify the relationship between time spent in the hold region and extensor burst duration, we isolated the extension region of the periodic orbit and performed a plane crossing analysis (**Fig. 2e,f**). Specifically, we constructed 2-D planes that were tangent to the 3-D trajectories at 7 locations along the extension region – 6 planes prior to the hold region and 1 after – and measured the time at which the single-cycle trajectories crossed each plane. Only trajectory crossing times for the plane after the hold region were highly predictive of the duration of extensor muscle activation across individual step cycles, precise on the scale of tens of milliseconds (*r*=0.95 and 0.82 for cat 1 and cat 2, respectively). This state space feature that we found in the extension region was distinctly different from what would be expected from a neural system where proportional changes in neural speed account for variations in the timing of motor output. In a proportional neural speed model, one might expect that variability in the generated extensor burst duration can be distributed over the time course of the trajectories evolving through the extension region of state space, however, we did not find this to be the case. To highlight how our results differed from this model, we measured the correlation between the observed plane crossing times and the expected crossing times from a proportional neural speed model. While the observed crossing times through the last plane along the arc (before entering the hold region) showed little correlation with the expected crossing times (cat 1: r=0.08, cat 2: r=0.17), the plane after the hold region showed strong correlation with the expected variation (cat 1: r=0.95, cat 2: r=0.74), highlighting how the encoding of burst duration variability was concentrated in the hold region.

Given the close correspondence between regions of interneuron state space and muscle output on individual step cycles, we reasoned that variability in state space trajectories should also be predictive of gross alterations of muscle output. Thus we next examined instances of *deletions*, where lapses in muscle activation happen spontaneously, sometimes disrupting the locomotor rhythm of either the ipsilateral antagonistic muscles or those on the contralateral side. Cat 2 exhibited multiple flexor deletion events, in which muscle activity broke from its stereotyped, periodic pattern (**Fig. 3a**). Prior to a deletion event (*pre-deletion*), extensor and flexor muscles activated in an expected alternating pattern. During a deletion, the extensor muscle remained active, while flexor activation was silenced. Following the deletion (*post-deletion*), extensor and flexor muscles returned to their previous alternating pattern of activation.

**Figure 3.**
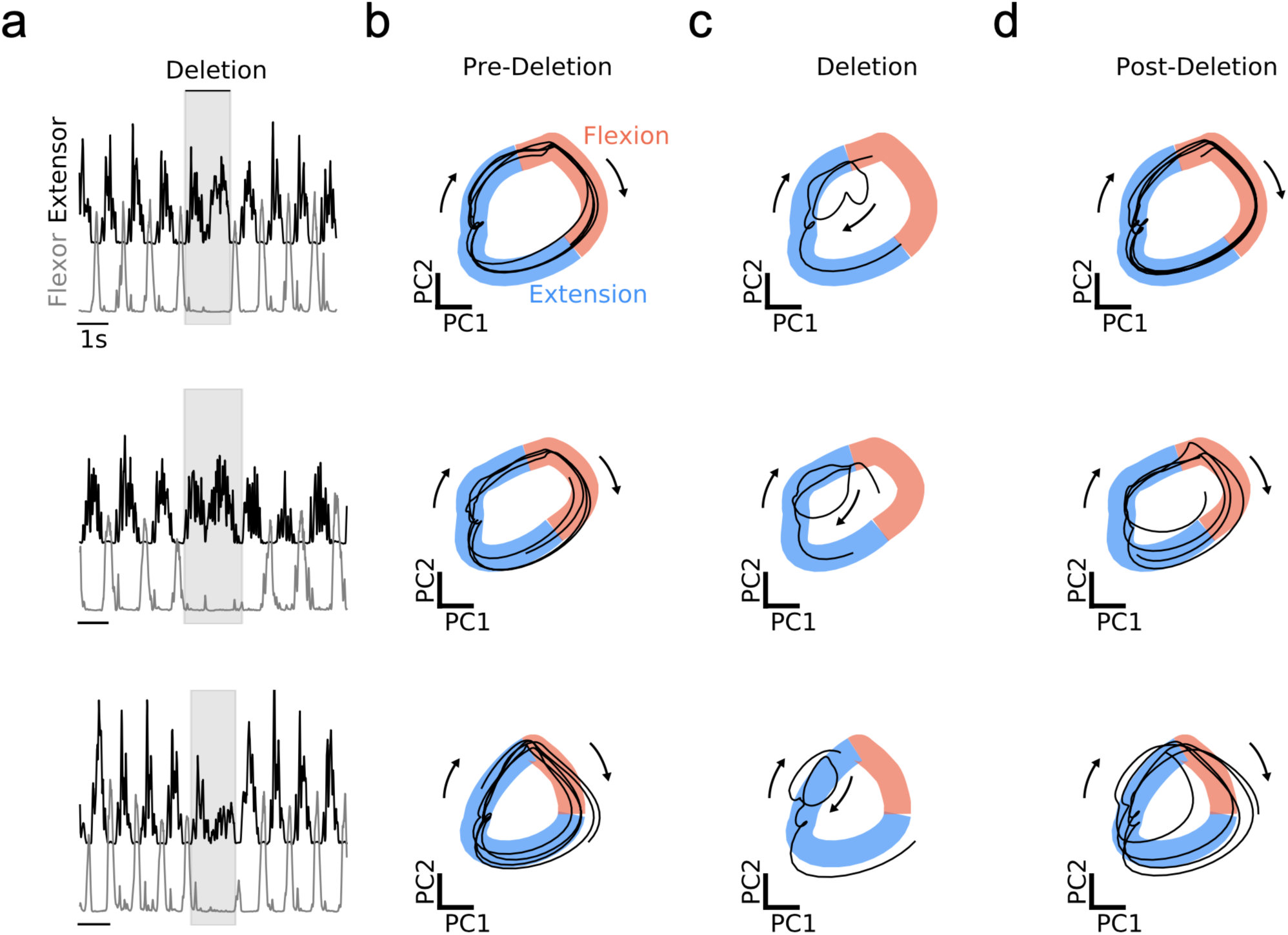
Aberrant muscle output corresponds with aberrant trajectories that avoid particular regions of interneuron state space. (a) Examples of muscle activity during three deletion events. Traces show ipsilateral extensor (black) and flexor (gray) muscle activations for multiple gait cycles. At the time of a deletion (shaded region), the extensor muscle remains active, while the flexor muscle activation is silenced. (b) Projection of the top two principal components of the spinal population activity during the pre-deletion period. State space coloring based on the flexion (red) and extension (blue) phases. (c) Same projection as in (b), during the deletion period. (d) Same projection as in (b), during the post-deletion period.

To determine whether deletions could be explained through the lens of interneuron state space, we visualized the top two principal components of interneuron population activity around the time of individual deletion events. While our previous analyses demonstrated that higher-order principal components were informative about muscle activations (**Fig. 1d)**, the top two principal components alone were sufficient to provide a simple visualization which revealed patterns that were readily interpretable. State space trajectories during pre-deletion cycles followed an expected repeating orbit that traversed the extensor and flexor regions (**Fig. 3b**). Prior to a flexor deletion, trajectories initially followed the same path through the extension region; however, rather than entering the flexion region, the deletion trajectories looped back into the extension region (**Fig. 3c**). After the deletion, state space trajectories returned to their consistent orbit (**Fig. 3d**). The avoidance of the flexion region during deletions is consistent with the idea that a regional organization of state space closely ties to the functional output of the interneuron network.

We next investigated which features of state space trajectories may be indicative of changes in the magnitude of muscle activations, which must be carefully adjusted by the spinal cord during each cycle. To maximize the precision of our analysis, we examined a brief time period just after extensor onset, which showed small (microvolt) differences in the magnitude of extensor activation across cycles (**Fig. 4a**). Simply coloring individual interneuron state space trajectories by the peak magnitude of extensor activation generated on each particular step revealed a close visual correspondence between the ordered deviations of trajectories and extensor muscle activation magnitude (**Fig. 4b**), which was also strongly correlated for both animals (**Fig. 4c**; *r*=-0.59 and -0.63 for cat 1 and cat 2, respectively). Previous models have suggested that increases in magnitude of motor output corresponded with an increased radius of rotation of the spinal population trajectories (Lindén et al., 2022). However, when we visualized planar projections of the top two principal components across both animals, the relationship between magnitude of muscle activation and rotation radius was inverted from predictions based on previous models (**Fig. 4d**). While these diverging observations may be attributed to a variety of factors (e.g., differences in model system, behavior, presence of sensory feedback, etc.), it highlights an important consideration whether rotations are a unifying organization principle underlying how spinal circuits regulate muscle activation magnitude.

**Figure 4.**
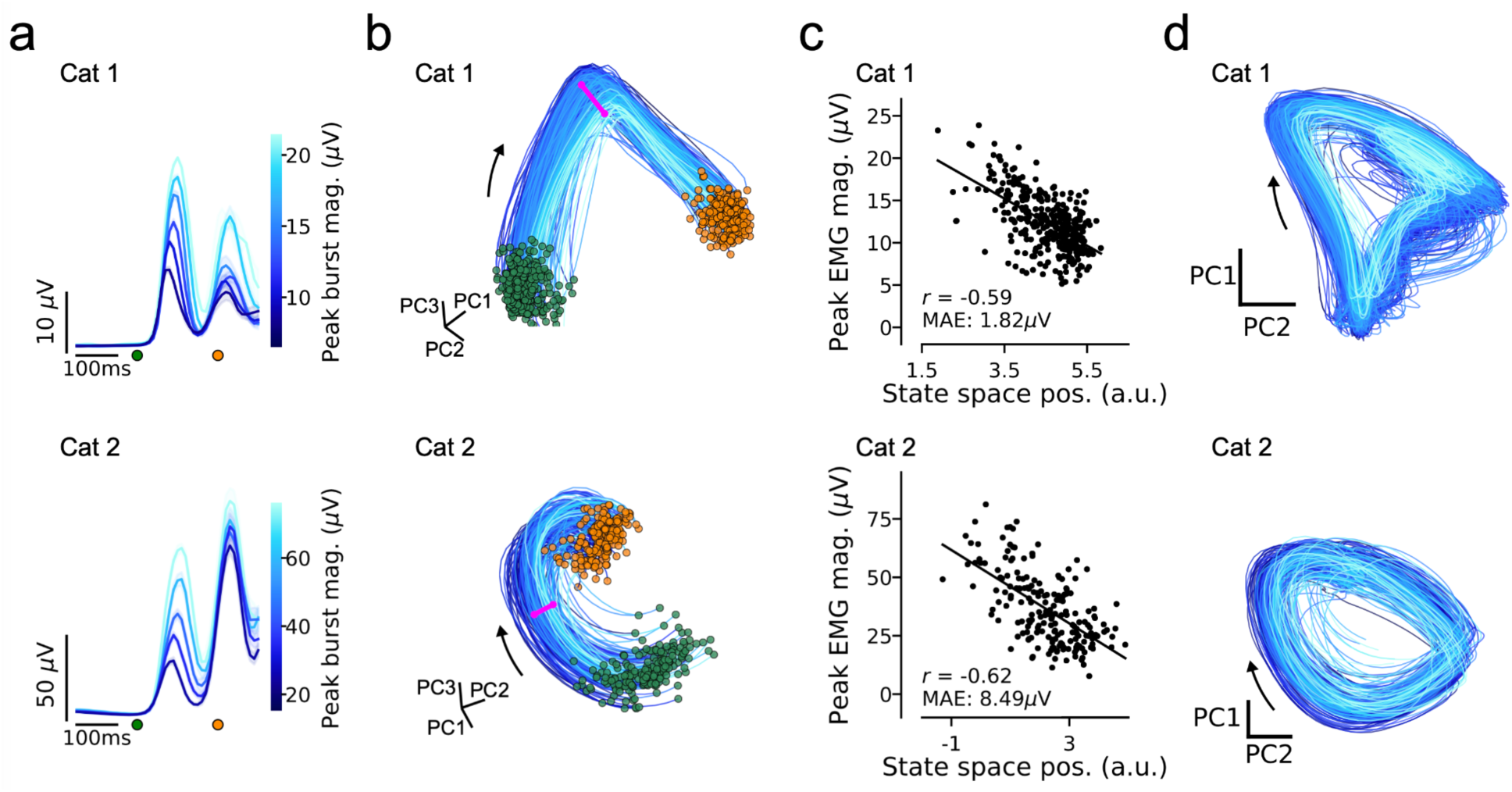
State space features of spinal interneuron population dynamics correlate with magnitude of muscle output. (a) Cycle-averaged ipsilateral extensor muscle (biceps anterior) activation around the time of extensor onset. Cycles were aligned to extensor onset (green dot) and grouped by the magnitude of extensor activation. Orange dot denotes 200ms after extensor onset. (b) Single cycle interneuron state space trajectories focused around extensor onset (200 ms). Trajectories for individual cycles are colored by extensor activation magnitude. Pink vector denotes the axis along which state space position was projected for computing correlations with extensor activation. Green and orange dots indicate the start (extensor onset) and end (200 ms after onset) of the trajectories, respectively. (c) Single-cycle relationship between extensor activation magnitude and interneuron state space position (cat 1: *r*=-0.59, median absolute error (MAE)=1.82 µV, cat 2: *r*=-0.62, MAE=8.49 µV). (d) 2-D projection of spinal interneuron activity onto top two principal components for individual cycles colored by extensor activation.

## Discussion

Our results establish that spinal interneuron state space – a level of abstraction relating individual interneurons and muscle activity – has identifiable features with a precise single-step correspondence to the duration and magnitude of muscle activations, on the scales of milliseconds and microvolts respectively. During air stepping, lumbar interneuron population trajectories traveled along an orbit where distinct state space regions corresponded with the flexion and extension phases of the step cycle. In interneuron state space, we found evidence of a localized hold region where the duration that state space trajectories remained in this region precisely corresponded with the duration of extensor activation with millisecond precision for individual steps. Further, we found that microvolt-scale adjustments in the magnitude of extensor muscle activation at the initiation of the step cycle corresponded with deviations in the path taken by population trajectories through interneuron state space. These findings indicate that using a state space framework to interpret lumbar population activity offers new insight into the computations that the spinal networks perform to shape motor output.

The flexibility to adjust locomotor pattern and speed on demand is attributed to the lumbar interneuronal network constituting the locomotor CPG (Frigon, 2012). Evidence from spinalized cats shows that the spinal cord, even in the absence of supraspinal input, retains the capacity to negotiate environmental challenges. For instance, spinalized cats maintain stable locomotion through adjusting the duration of the extension phase in response to changes in treadmill speed (Barbeau and Rossignol, 1987; Forssberg et al., 1980; Frigon et al., 2013; Grillner and Wallén, 1985; Lovely et al., 1990) and during stumbling corrective responses (Forssberg et al., 1977). In both cats (Halbertsma, 1983) and humans (Grillner et al., 1979), the period of individual step cycles is primarily determined by the duration of extension during the stance phase, whereas the swing phase remains relatively stable. We similarly found that variations in the duration of the extension phase of air stepping mainly accounted for single-cycle adjustments to locomotor speed. The extension phase was compactly described by two distinct state space features: 1) a large arc through state space at the initiation of extension and 2) a hold region that corresponded with varying durations of extension.

The contrast between the way interneuron trajectories evolve during the initiation and hold period of extensor activation may indicate differences in underlying neural mechanisms governing these two operations. Analyzing the consistency with which single-cycle population trajectories evolve through a given region of state space has been shown to be indicative of the type of dynamics (i.e., smooth, autonomous vs input-driven) that govern a given operation performed by the neural population (Russo et al., 2018). For example, during planned cycling movements, motor cortical population trajectories smoothly flow in a similar direction and speed through specific regions of state space. In contrast, during the same behavior, population trajectories in the proprioceptive region of somatosensory cortex are entangled in state space and do not show clear patterns of how trajectories should predictably evolve with respect to speed or direction. While the ordered, predictable evolution of the interneuron population trajectories through state space at extensor initiation is more consistent with a smooth, autonomous response, the confined and clashing trajectories in the hold region are more indicative of an input-driven response. In the spinal cat, where supraspinal inputs from upstream brain areas are absent, sensory afferents signaling stretch and load of the extended hindlimb are plausible feedback signals that may confine state space trajectories to the hold region, thereby extending the duration of the extension phase (Frigon et al., 2021). The activation of hindlimb group 1 afferents is known to prolong the stance phase of the current step cycle, consequently delaying onset of the subsequent swing phase during fictive (Angel et al., 1996; Conway et al., 1987; Gossard et al., 1994) and treadmill locomotion (Pearson and Collins, 1993; Whelan et al., 1995) in spinalized, decerebrate cats. Future studies that are able to directly probe and manipulate the activity of hindlimb afferents in an experimentally-controlled fashion can further determine their roles in generating the hold region we found.

While we identified that the hold region was an important state space feature in governing the duration of extensor burst during air stepping, we acknowledge that the utility of this feature may not be apparent or implemented across all behaviors. Behaviors that are dominated by flexor-muscle activity or require more variation in the duration of the swing phase (e.g., obstacle avoidance, (Lecomte et al., 2023)) may require different mechanisms to generate that type of motor output variability that we did not observe in the spinal cat during air-stepping. Additionally, the extension bias we observed may also be attributed in part due to the experimental conditions (i.e., spinalized animal, clonidine-induced, air stepping with no ground reaction forces). Recordings during locomotion in intact animals may reveal additional state space features that underlie more natural locomotor behaviors. Conversely, fictive locomotor preparations, where the spinal cord is producing locomotor patterns in the absence of sensory input, could provide a contrasting insight into the intrinsic capacity of interneuronal networks to generate locomotion. Understanding how state space features such as the hold region are employed across multiple behaviors will clarify how conserved or distinct these operating modes are within interneuron state space.

Research into the biological implementation of the locomotor CPG has largely focused on interconnections between populations of spinal interneurons, ranging from the earliest “half-center” circuit models that proposed interactions between populations of excitatory and inhibitory interneurons (Brown, 1911), to recent models that integrate genetically-identified interneuron subtypes (Rybak et al., 2015). Several of these proposed models can reproduce observed locomotor activity patterns to varying degrees (McCrea and Rybak, 2008). However, the problem of multiple realizability (i.e., different hypothesized connectivity patterns and posited neural populations can reproduce muscle activity patterns with similar accuracy), and the difficulty of linking the activity of the simulated interneuron populations to actual electrophysiological recordings, makes it challenging to know which putative model is best. Our data-driven approach to directly uncover dominant patterns of interneuron population activity complements such circuit modeling approaches. For example, identified state space features could serve as new targets or constraints for developing circuit models. Additional studies that characterize the effects of cell type-specific or neuromodulatory perturbations on state space features will further serve to constrain circuit models.

A primary objective in both clinical management and the development of therapies for spinal cord injury (SCI) is to re-engage the locomotor CPG to regain functional capabilities post-injury (Grillner, 2002). Currently, assessment of the effectiveness of experimental therapies relies solely on evaluating gross motor functions (e.g., observed muscle activity), without detailed insights into the actual operations of the spinal cord network that governs this motor output. Thus, there is a significant need for tools that precisely identify the operational status of the spinal locomotor circuitry to inform research related to SCI recovery. Given the presence of similar state space features in multiple subjects, spinal interneuron state space may be a suitable framework for gauging network operations in the cord, evaluating spinal network changes following SCI and other diseases, and serving as a physiological target for guiding functional restoration approaches such as spinal cord stimulation (Barra et al., 2022; Capogrosso et al., 2016; Harkema et al., 2011; Wagner et al., 2018) or pharmacological interventions (Rabchevsky et al., 2011; Song et al., 2019).

A large body of literature, developed over a century, provides a wealth of detail on spinal cord anatomy and cell type distributions (Roome and Levine, 2023; Sengupta and Bagnall, 2023). Emerging technologies for large-scale, chronic recording and stimulation of spinal populations (Berg et al., 2009; Metuh et al., 2024; Wu et al., 2023), as well as the advancements in approaches to monitor and manipulate the well-defined spinal inputs (i.e., tactile, proprioceptive, nociceptive neurons) and outputs (i.e., motorneurons (Chung et al., 2023)) are rapidly being realized. This unique access to both the inputs, outputs, and internal state of a neural system make the spinal cord a unique testbed to apply state space approaches to uncover the neural principles underlying flexible pattern generation with unprecedented precision and explainability.

## Acknowledgements

We thank Drs. Chantal McMahon and Michel Lemay for collecting and sharing the data. We also thank Toossi and colleagues for allowing re-use of their cat spinal cord schematic for Figs 1 & 2 in this paper (Toossi et al., 2021). We thank Ariel Levine, Andrew Pruszynski, Chris Rodgers, and Saurabh Vyas for feedback on the manuscript. This work was supported by NSF NCS 1835364, the Alfred P. Sloan Foundation, NIH BRAIN/NIDA RF1 DA055667, NIH-NINDS/OD DP2NS127291, the Simons Foundation as part of the Simons-Emory International Consortium on Motor Control (CP), Air Force Office of Scientific Research FA9550-23-1-0727 (CP, NAY), and the Emory Neurosurgery Catalyst Grant (NAY).

## Data availability statement

Data will be made publicly available upon publication.

## Supplementary Figures

**Supplementary Figure 1.**
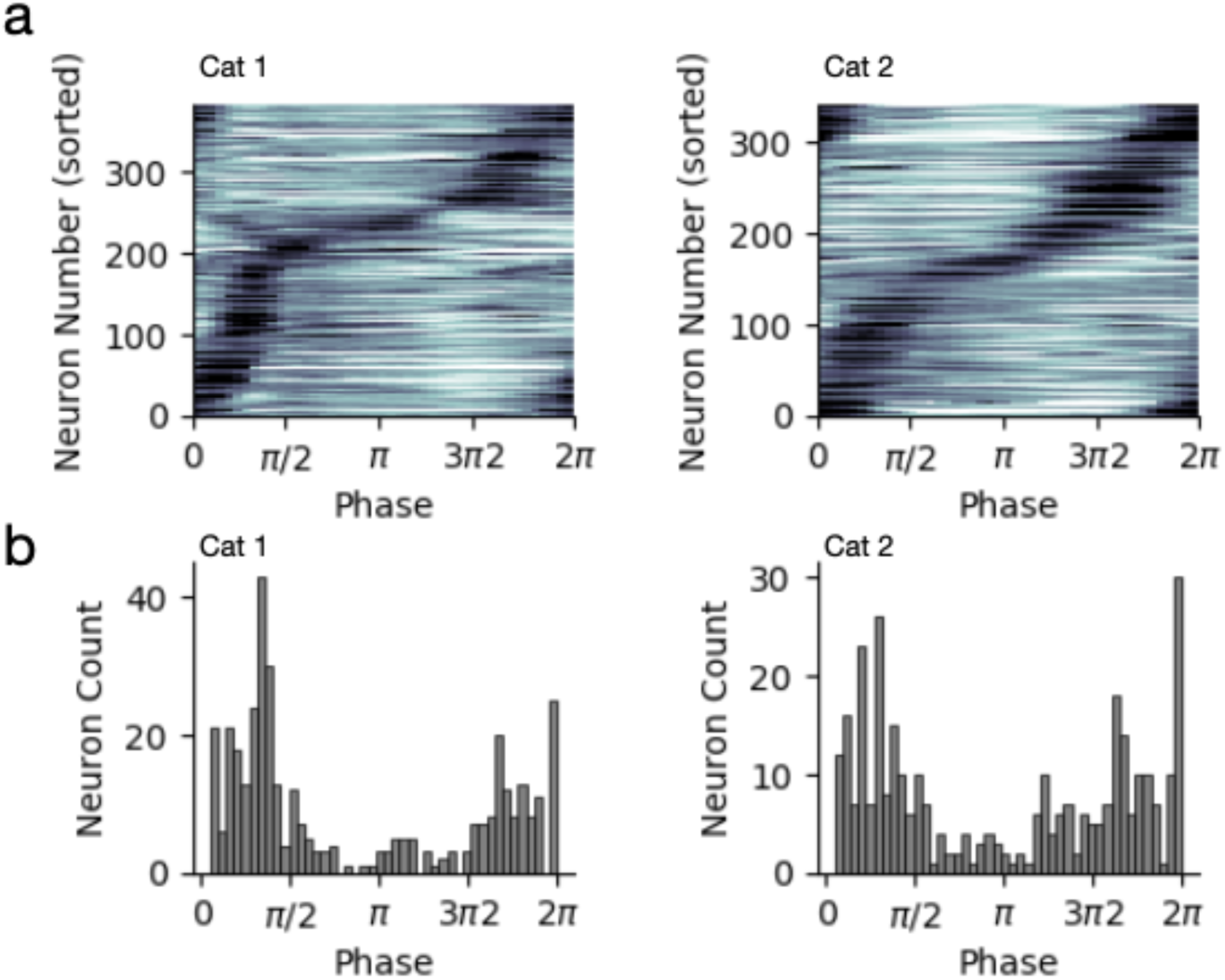
Interneuron population activity normalized to step phase. (a) Interneuron population activity combined across sessions (Cat 1: 382 neurons, 341 neurons) sorted based on the peak firing rates of the population in order of increasing phase. (b) Phase distribution of the peak firing rates of the interneuron population.

**Supplementary Figure 2.**
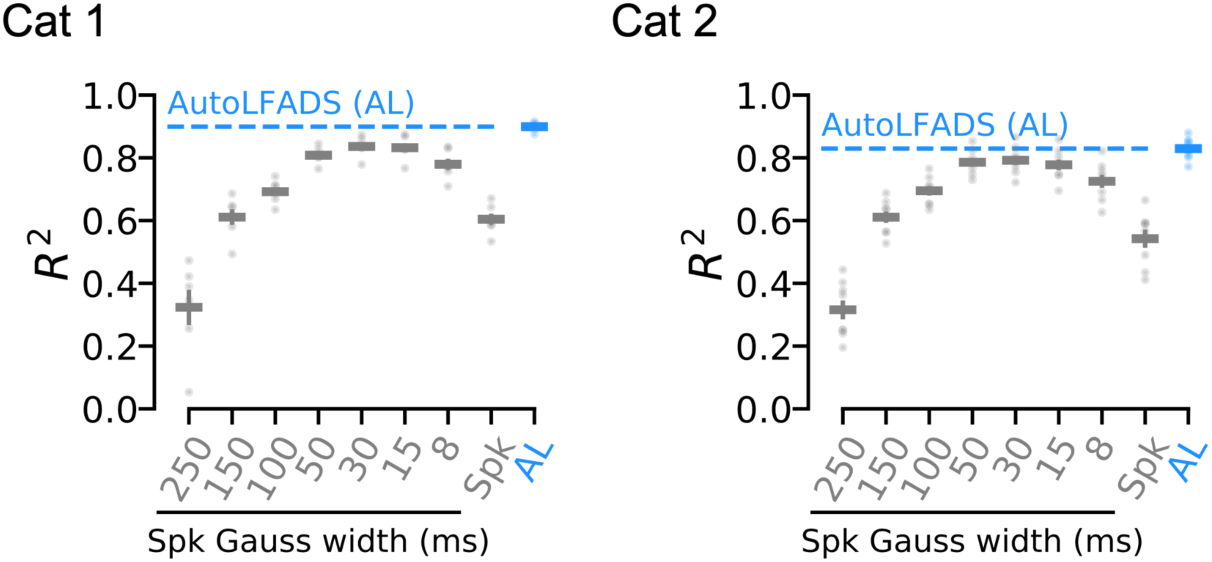
Decoding muscle activations from different estimates of neural firing rates. Comparison of predictions of muscle activations using single-timepoint optimal linear estimation from different estimates of spinal firing rates. Predictions from AutoLFADS-inferred firing rates outperformed predictions from Gaussian-smoothed firing rate estimates with kernels of various widths (quantified using R^2). Small dots show prediction accuracy for a given recording session, summarized by the horizontal line representing average decoding accuracy across sessions (cat1: 6 sessions, cat2: 8 sessions) for a given approach.

**Supplementary Figure 3.**
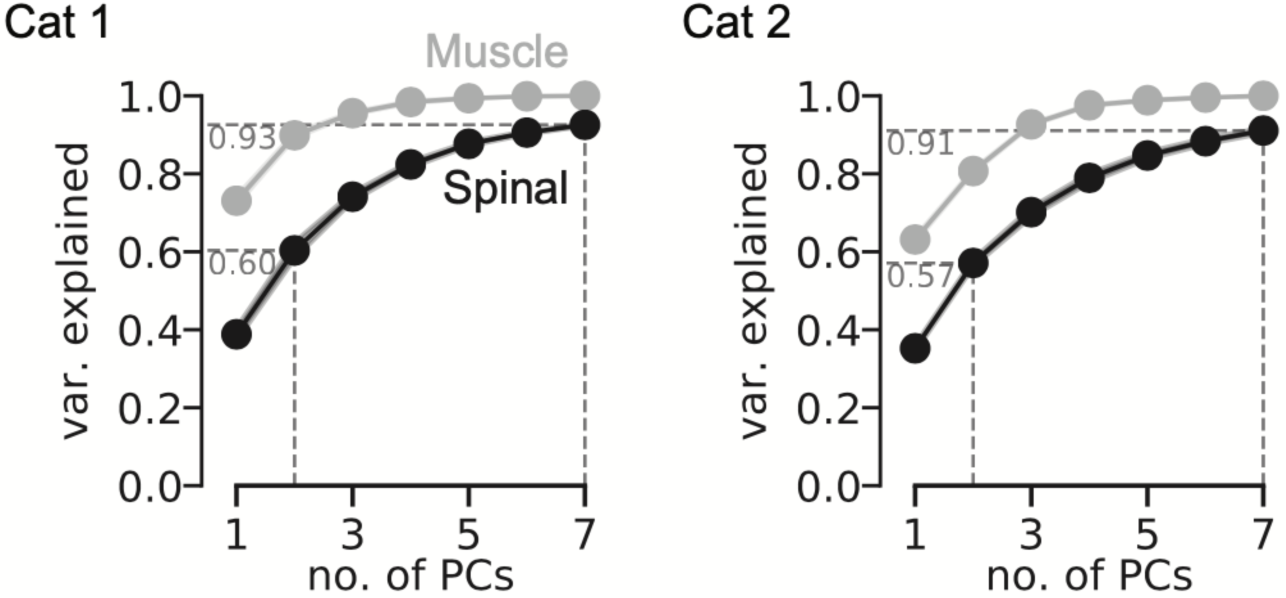
Comparison of variance explained using PCA. Cumulative variance explained with increasing number of principal components for ipsilateral muscle activations (gray) and spinal population activity (black). PCA was applied individually to each session’s activity. Mean and SEM across estimates of variance explained for individual sessions shown.

**Supplementary Figure 4.**
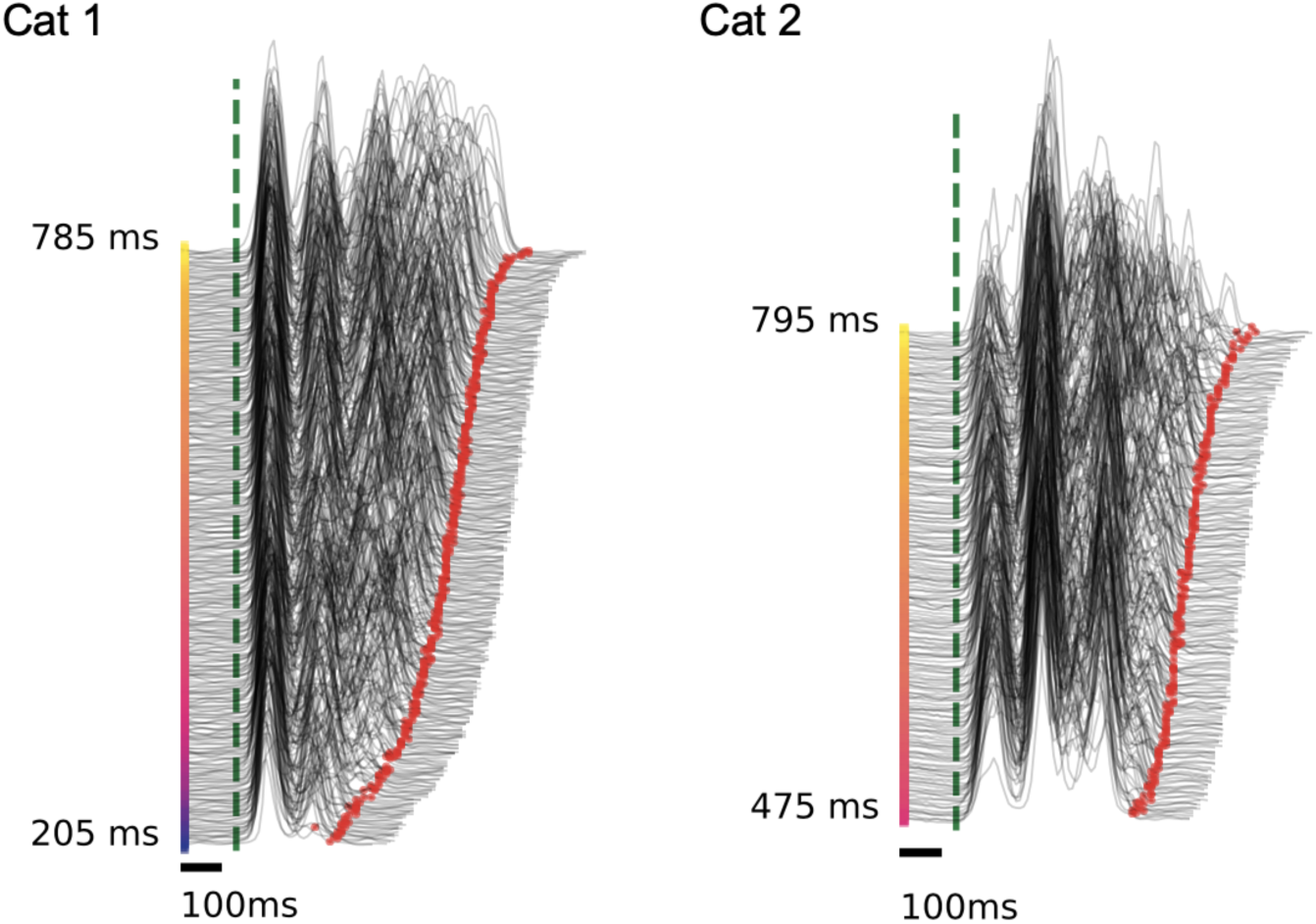
Identification of onsets and offsets of extensor burst. Single-cycle muscle activation traces for an extensor muscle (biceps femoris anterior). Muscle activations were aligned to the identified onset (green dashed line), marked with the identified offset (red dot), and ordered based on the duration of extension (cat 1: 335 cycles, cat 2: 223 cycles).

**Supplementary Video 1.**
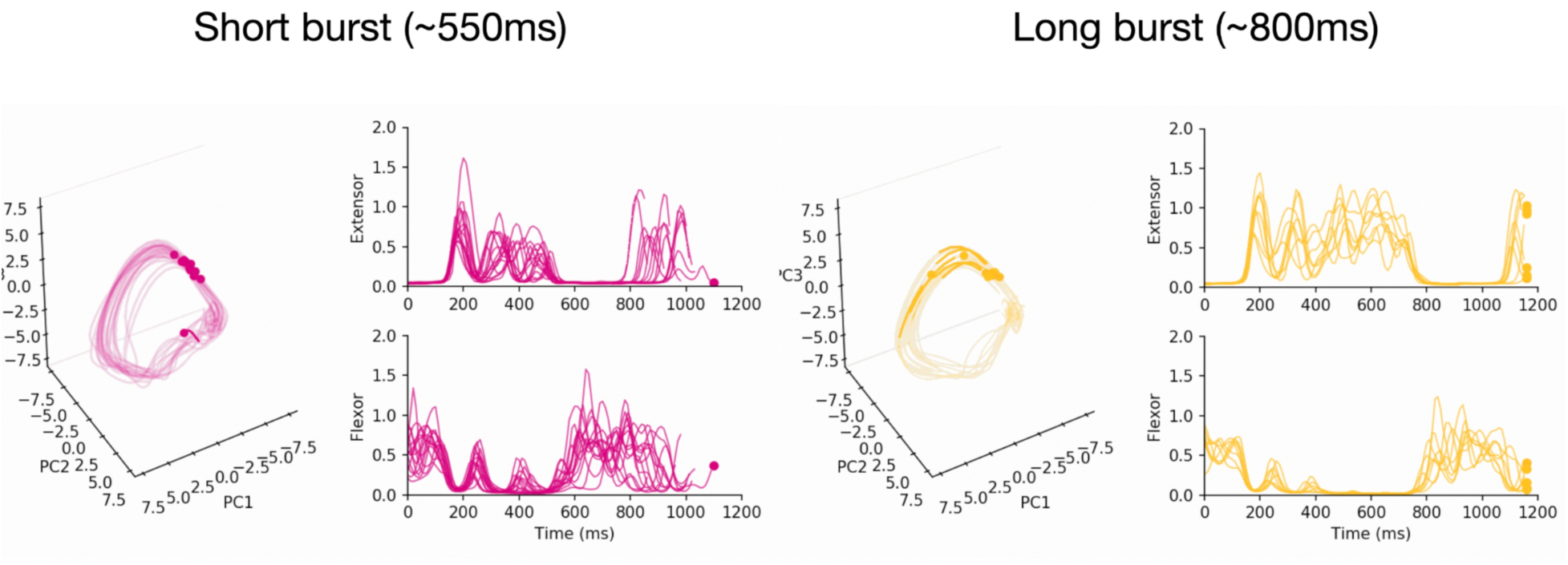
Link: https://drive.google.com/file/d/1av_fo7qB9RvRgQLyXfHxksV6AzVnBTY3/view?usp=sharing

## Methods

### Datasets

#### Experimental setup

We analyzed previously-collected datasets from an air-stepping model in sub-acute spinalized cats whose data collection procedures have been presented in detail previously (McMahon et al., 2022). Briefly: data were collected from two adult domestic shorthair female cats. First, a T11/T12 spinal transection was performed 22-23 days before the recording session. No locomotor training was provided to the animals at any point before the terminal experiment. On the day of the terminal recording experiment, three sets of surgical procedures were performed on the anesthetized animal: 1) a laminectomy to expose the lumbar cord, 2) implantation of bipolar EMG electrodes into seven muscles of each hindlimb to record muscle activity, and 3) a mid-collicular decerebration to discontinue anesthesia. Following the surgical procedures, animals were transferred to a stereotaxic frame where the spinal vertebrae were securely clamped to the frame. At approximately one hour after decerebration, clonidine (500 µg/kg) was administered intravenously to prime air-stepping. Experimental recording sessions began once air stepping was inducible through perineal stimulation.

#### Neural recordings

Spinal interneuron activity was simultaneously recorded using two 64-site microelectrode arrays (model A8x8-5mm-200-200-177, NeuroNexus, Ann Arbor, MI) implanted on the right side of the spinal cord. Multiple recording sessions were collected with each animal, where between some recording sessions, one of the arrays was moved to sample spinal interneuron activity at different rostral/caudal locations between L4 and L7. Arrays were implanted at or near the dorsal root entry zone to an approximate depth of 3000 microns, such that the electrode contacts covered a range from 1500 to 3000 microns deep, placing them in the intermediate zone of the spinal cord between the dorsal and ventral horns. This was done intentionally so that the recorded units were spinal interneurons and not motorneurons. Recorded multi-electrode activity was spike sorted using UltaMegaSort2000 (Joshua et al., 2007). The sorted single-unit activity met the following criteria: 1) greater than 5 spikes per step for at least 5 consecutive steps, 2) greater than 200 spikes in a session, 3) contained less than 1.5% refractory period violations for a refractory period of 1.5ms, 4) signal-to-noise ratio greater than 1.5, and 5) did not respond antidromically to sciatic nerve stimulation as indicative of motor neurons. Neural activity was collected during a total of 13 and 27 air-stepping sessions for Cat 1 and Cat 2, respectively. Each air-stepping session consisted of a ∼5 s rest period, followed by ∼50-70 s of air stepping, and completed with another ∼5 s rest period; however, the quality of air-stepping, as well as the neuronal yield, varied from session-to-session. Given this variability we sub-selected sessions for analysis based on the presence of consistent air-stepping throughout the entire duration of the session and high neuronal yield. Neuronal yield was generally higher for Cat 1 than Cat 2, so we set the thresholds for the minimum number of neurons recorded in session accordingly (cat 1: >55 neurons, cat 2: >35 neurons). Following the sub-selection process, we analyzed 6 and 8 air-stepping sessions from Cat 1 (382 neurons, 335 step cycles) and Cat 2 (341 neurons, 193 step cycles), respectively.

**Table 1.** Cat 1 Dataset Information.

| Number of Neurons | Array 1 Location | Array 2 Location | Length of recording (s) |
| --- | --- | --- | --- |
| 71 | L3 | L6 | 75.3 |
| 62 | L3 | L5 | 81.6 |
| 65 | L3 | L5 | 83.3 |
| 63 | L3 | L5 | 77.4 |
| 62 | L3 | L5 | 77.9 |
| 59 | L3 | L5 | 67.4 |

**Table 2.**
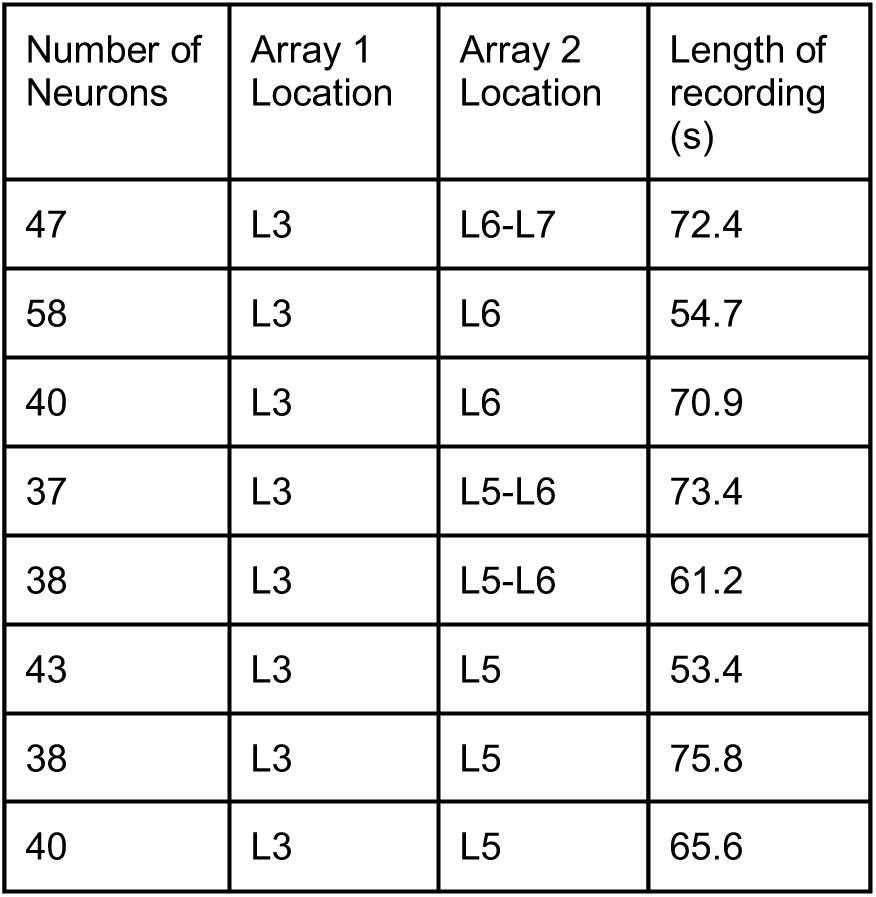
Cat 1 Dataset Information.

#### EMG recordings

To monitor muscle output, intramuscular EMG was recorded from seven muscles spanning each hindlimb, including three flexors (sartorius anterior, biceps femoris posterior, and tibialis anterior) and four extensors (biceps femoris anterior, vastus lateralis, gastrocnemius medialis, and soleus). Each muscle was implanted with bifilar electrodes constructed with insulated multi-strand stainless steel wires (AS 633; Cooner Wire, Chatsworth, CA). Electrodes were implanted into the belly of the muscle and secured onto the fascia with sutures. Electrode placement was verified by stimulation of the electrode and observation of the resulting muscle’s twitches.

### Modeling

#### Applying AutoLFADS to de-noise neural recordings

We applied the deep learning technique AutoLFADS to de-noise recorded neural activity to obtain high-fidelity estimates of interneuron population firing rates and muscle activations (Keshtkaran et al., 2022; Wimalasena et al., 2022). Briefly, AutoLFADS models the spatial and temporal regularities underlying neural recordings using recurrent neural networks (RNN). Data is passed into the LFADS model through a set of bi-directional RNN encoders that estimate an initial state of one RNN (termed the *Generator*) and a set of time-varying encoding states that are passed into a second RNN (termed the *Controller*). The Generator is run forward in time from the initial state, where at each timestep, time-varying inputs estimated by the Controller are applied to the Generator state, simulating a non-autonomous dynamical system. The output of the Generator is then passed through two successive linear read-out matrices. The first linear read-out matrix with weights *W^fac^* limits the output (termed the *output factors*) of the Generator to describe the patterns of the observed neural activity with a fewer number of dimensions. The output factors are then passed through a second linear read-out matrix with weights *W^rate^*, followed by an exponential nonlinearity, to map back to the original data, where a reconstruction cost (negative log-likelihood) is computed between the estimates of the LFADS model and the original data and used to train the model via backpropagation. AutoLFADS trains multiple LFADS models (with different regularization hyperparameters) in parallel and selects the best performing models (lowest validation cost) via an automated hyperparameter optimization called population-based training (PBT, (Jaderberg et al., 2017)).

To leverage neural recordings collected across multiple recording sessions, we combined the automated hyperparameter optimization of AutoLFADS (Keshtkaran et al., 2022; Keshtkaran and Pandarinath, 2019) with *stitching* (Pandarinath et al., 2018). In a stitched LFADS model, data from a given recording session is passed through a pre-initialized *read-in* matrix that linearly maps recorded neural activity onto a set of *input factors*. Each session-specific read-in matrix is fixed with weights *W^input^*, which are learned from principal components regression (PCR). To perform PCR, we cycle-averaged data in a window around extensor onset (100ms prior, 600ms after) for each session. Cycle-averages were concatenated across sessions such that the channels for each session were considered as independent input features (termed *global cycle-averages*). We then applied PCA to the mean-centered global cycle-averages and sub-selected the top *k* principal components (PCs) to serve as our *global PCs* (k=10 for EMG, k=20 for spikes). We then learn the weights *W^input^*^’^ by fitting a Ridge regressor between the mean-centered cycle-averaged data to the global PCs (L2=1e-2). The bias term of affine transformation was obtained by passing the means of the cycle-averaged data through the learned *W^input^*.

The input factors from all sessions are passed through a shared LFADS model to produce generator states as described above, followed by a shared *W^fac^* to obtain output factors within a common latent space across all sessions. The output factors are then passed through session-specific read-out matrix *W^rate^* that is randomly initialized and learned during model training.

We performed two separate applications of AutoLFADS: 1) to infer de-noised spinal interneuron firing rates from sorted spiking activity and 2) to infer high-fidelity muscle activation estimates from rectified EMG signals (Wimalasena et al., 2022). All fixed hyperparameters (HPs) used for modeling can be found in Table 3. To briefly summarize, most model hyperparameters were consistent across LFADS models applied to muscle and interneuron data. The differences (highlighted at the top of Table 3) were in some of the dimensionalities of the internal LFADS representations (e.g., inferred inputs, input/output factors) and the reconstruction cost computed during training (i.e., Gamma or Poisson log-likelihood). To prevent overfitting to individual spikes or overfitting to correlated noise across EMG channels, we applied data augmentation strategies *temporal jitter* (applied to discrete binned spike counts) and *temporal shift* (applied to continuous EMG) during model training. Temporal jitter involves shifting individual spikes randomly in time, up to *k* bins (a settable HP) before or after their original time bin prior to being input to the LFADS model. Temporal shift involves applying random temporal offsets of *k* bins (a settable HP) separately for each channel during model training. Data input to the model encoders and the data used to compute reconstruction were shifted differently to prevent the network from overfitting by drawing a direct correspondence between specific values of the input and output. For spiking and EMG models, we used a max temporal jitter of 1 bin and a max temporal shift of 1 bin, respectively.

All HPs searched using the AutoLFADS framework can be found in Table 4. To briefly summarize, PBT was allowed to optimize the learning rate and five regularization HPs: the L2 penalties for the Generator and Controller, the scaling factors for the KL penalties applied to the initial conditions and the controller outputs (i.e., the time-varying inputs to the Generator), the coordinated dropout rate (Keshtkaran and Pandarinath, 2019), and the dropout rate. At the beginning of training, all models began with a learning rate of 4e-3 and a coordinated dropout rate of 0.5, since varying these parameters can have an effect on how fast the model trains. We trained 18 LFADS models in parallel (with different hyperparameter combinations) for 10 epochs (one generation) before PBT performed a selection process to choose higher performing models and eliminate models with poor performance. All parameters for PBT are summarized in Table 5.

**Table 4.**
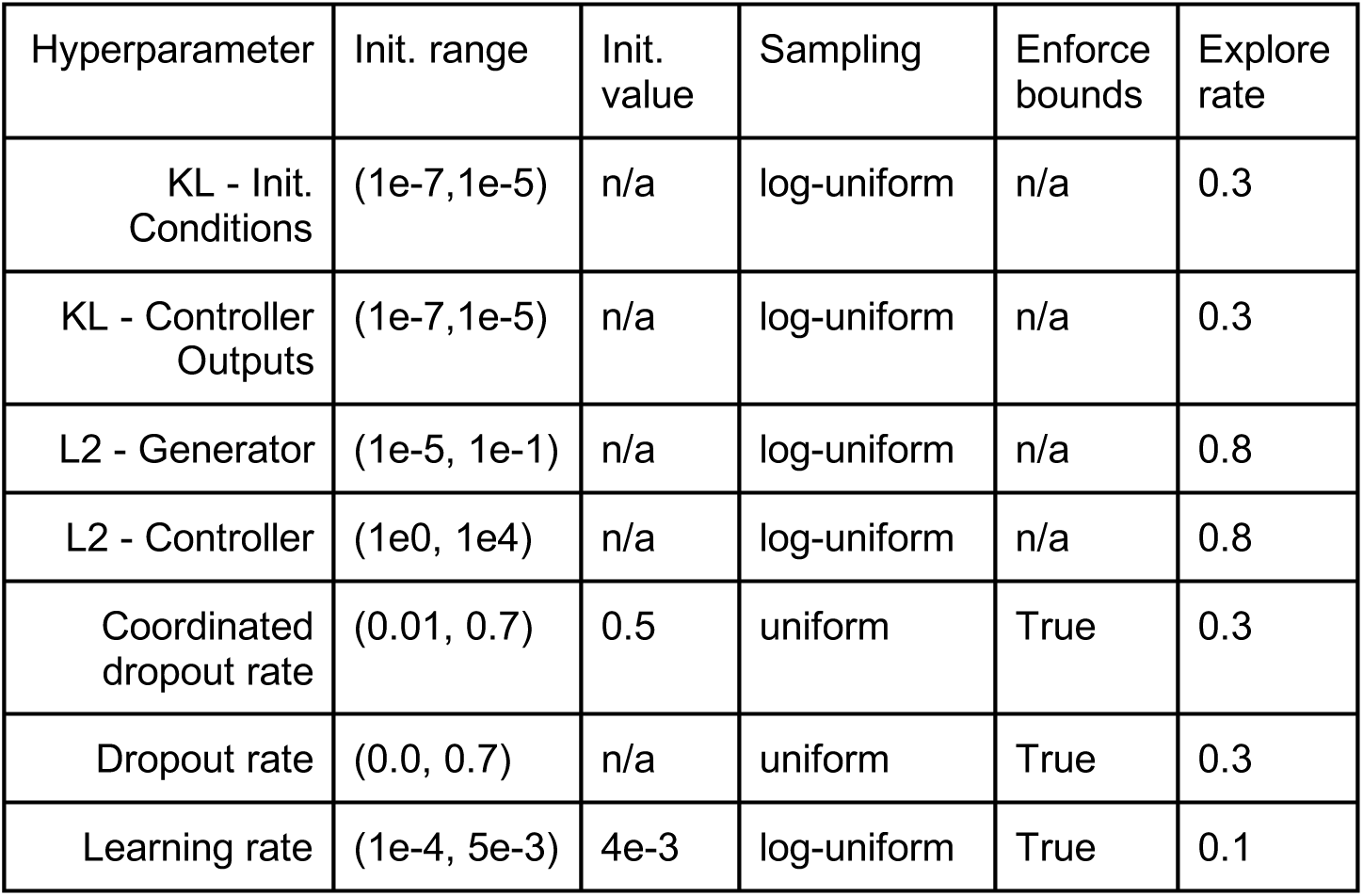
LFADS hyperparameters search ranges and initializations for PBT.

| Hyperparameter | Init. range | Init. value | Sampling | Enforce bounds | Explore rate |
| --- | --- | --- | --- | --- | --- |
| KL - Init. Conditions | (1e-7,1e-5) | n/a | log-uniform | n/a | 0.3 |
| KL - Controller Outputs | (1e-7,1e-5) | n/a | log-uniform | n/a | 0.3 |
| L2 - Generator | (1e-5, 1e-1) | n/a | log-uniform | n/a | 0.8 |
| L2 - Controller | (1e0, 1e4) | n/a | log-uniform | n/a | 0.8 |
| Coordinated dropout rate | (0.01, 0.7) | 0.5 | uniform | True | 0.3 |
| Dropout rate | (0.0, 0.7) | n/a | uniform | True | 0.3 |
| Learning rate | (1e-4, 5e-3) | 4e-3 | log-uniform | True | 0.1 |

**Table 5.** Population Based Training (PBT) parameters.

| Hyperparameter | Value |
| --- | --- |
| # of epochs per generation | 10 |
| Max # of generations | 200 |
| Patience (generations) | 15 |
| Evaluation Metric | Smoothed validation recon. cost |
| # of workers per generation | 18 |

To prepare data for modeling, we performed a set of pre-processing steps for the EMG and spiking data. To prepare the spiking data, we performed a pairwise correlation analysis between the 1-ms binned spiking activity of all neurons in a given session. Pairs of neurons with a correlation greater than or equal to 0.1 were identified and were passed to an iterative process that removed individual neurons from the dataset until all correlations between include neurons were below the set threshold. To prepare the EMG data, raw recordings were first filtered using a 4th order Butterworth high-pass filter at 65 Hz to reduce movement artifacts and suppress any powerline noise (60 Hz), followed by full-wave rectification and resampling to 1000 Hz. To remove artifacts and prepare multi-channel EMG data for LFADS modeling, we first clipped each channel’s activity such that any activation above a threshold (99th percentile of channel’s activation) was set to the threshold. We then normalized each channel’s activity by the 95th percentile so that all channels had comparable variance during modeling. To prepare the training and validation datasets used for modeling, Spiking data was re-binned to 10-ms bins and EMG data was resampled to 100 Hz and re-rectified to ensure data was strictly positive. Continuous data from each session were segmented into 1-s (100 bins) windows that were overlapped by 200-ms. After applying this windowing procedure to the entire dataset, we separated every fifth consecutive window (20% of all windows) to serve as the validation set. The remaining windows were used for training.

**Table 3.** Fixed LFADS model hyperparameters for EMG and spikes models.

| Hyperparameter | EMG | Spikes |
| --- | --- | --- |
| Recon. cost | Gamma | Poisson |
| Input factor dim. | 10 | 20 |
| Output factor dim. | 10 | 20 |
| Inferred input dim. | 3 | 6 |
| Initial condition dim. | 20 | 20 |
| Generator dim. | 100 | 100 |
| Controller dim. | 100 | 100 |
| Initial condition encoder dim. | 100 | 100 |
| Controller input encoder dim. | 100 | 100 |
| Batch size | 128 | 128 |
| Sequence length (bins) | 100 | 100 |
| Bin size (ms) | 10 | 10 |
| Temporal shift (bins) | 1 | N/A |
| Temporal jitter (bins) | N/A | 1 |
| L2/KL penalty ramp (epochs) | 10 | 10 |
| Max gradient norm | 200 | 200 |
| Adam opt. epsilon | 1e-8 | 1e-8 |

### Analysis

#### Computing onset/offset times

Extensor onset and offset times were initially computed on the de-noised muscle activation trace (i.e., output of AutoLFADS-EMG model) of the right biceps femoris anterior using a custom detection algorithm we developed. The detection algorithm analyzed the differentiated muscle activation trace to identify event timepoints where muscle activation rapidly increased or decreased. These initial estimates of extensor onset were used to align data to find low-dimensional state space descriptions of spinal interneuron population activity and the multi-muscle activations across recording sessions using principal components regression (PCR, see *Finding shared low-dimensional state-space descriptions*). After identifying shared low-dimensional state space descriptions of the right leg muscle activations using PCR, we measured the events where activity along the top PC crossed a baseline value. This transition within the multi-muscle activation state space closely corresponded with the timings of when the flexor/extensor onset/offset times, confirmed visually from the aligned muscle activation traces across sessions (e.g. Fig. 2a, Supp. Fig. 2).

#### Finding shared low-dimensional state-space descriptions

Following AutoLFADS modeling, we applied PCR to 1) further identify shared components across multiple recording sessions, and 2) organize components based on their total contributions to the overall variance in the AutoLFADS output. To fit PCR, we followed the same process described above to fit the read-in matrices for AutoLFADS modeling, however we used different time windows for the neural and muscle activity. We fit PCR for neural activity on a smaller window around extensor onset (AutoLFADS factors and smoothed spikes: (60ms prior, 100ms after)), and used the larger window for muscle activations (300ms prior, 400ms after), enabling us to fit PCR on a window that included times when both flexor and extensor muscles were maximally active. When fitting the read-in matrix for each session, we used an L2 regularization coefficient of 1e1 for the AutoLFADS factors (from spikes model) and the AutoLFADS-inferred muscle activations and a regularization coefficient of 1e-2 for smoothed spiking activity.

#### Selection of cycles for analysis

We aimed to analyze spinal interneuron population dynamics during periods of normal gait (excluding the deletion analysis) where we could easily identify the onset/offset timings of flexor and extensor muscle activations. To do so, we limited our analysis to cycles with extension phases within 200- to 800-ms and flexion phases between 200- and 460-ms for Cat 1, and 200- to 510-ms for Cat 2. The range of flexion phase included for analysis was increased to capture more cycles for Cat 2 where we observed stable locomotion, however, there were some cycles where calculated flexor onset and offset timings were imprecise, thus we excluded them (Cat 1: 4 cycles, Cat 2: 18 cycles). Additionally, to exclude periods in which locomotion was starting or stopping, we excluded the first and last three cycles from each session. We also excluded cycles around periods when air-stepping paused for a brief moment during a trial (Cat 1: 2 cycles, Cat 2: 5 cycles).

#### Computing attributes of muscle activity

To compute burst magnitude, we took the peak value of the de-noised muscle activation estimate in a window aligned to extensor onset (80ms after, 120ms after). To compute extensor burst duration, we computed the difference between the extensor offset and onset times we identified.

#### Relating interneuron state space to burst duration

We computed a vector to establish a 2D plane within the state space for the purpose of measuring the timing of the spinal interneuron state as it crossed through this plane. We wanted to appropriately place this plane to capture the evolution of activity around the extensor burst offset event. To compute this vector, we determined the average 3D state space positions at two timepoints relative to the extensor burst offset (*point_1*: 100ms prior, *point_2*: 40ms prior). Using *point_1* and *point_2*, we constructed a vector that pointed in the dominant direction of flow of the state space trajectories across all cycles. We then projected the 3D spinal interneuron state space trajectories onto this vector. We measured the zero-crossing time along this projected dimension, which indicated the moment when the state space trajectory intersects the plane orthogonal to the vector. The timing of these events served as our identified interneuron state space crossings.

#### Relating interneuron state space to burst magnitude

We computed a second vector to span the direction in spinal interneuron state space that captured the muscle activation magnitude encoding in the spinal interneuron state space trajectories. To do this, we isolated state space trajectories aligned to the timing of extensor onset. We then separated the trajectories into two groups of ten cycles around the 50th and 95th quantile of measured extensor burst magnitudes, respectively. For each group, we computed the average 3D state space position within a 20ms window around a specific timepoint after extensor burst onset (*point_1*: 70ms and 80ms, *point_2*: 70ms and 100ms for Cat 1 and Cat 2, respectively). Using *point_1* and *point_2*, we constructed a vector and projected the 3D spinal interneuron state space trajectories onto this dimension. We visualized the trajectories along this projected dimension, colored by extensor burst magnitude, and identified a timepoint near extensor onset that showed clear ordering of the trajectories based on burst magnitude (Cat 1: 70ms, Cat 2: 40ms). The state space positions at these identified timepoints were used to compute linear correlations with the extensor burst magnitude.

#### Linear decoding

We performed zero-lag single-timepoint optimal linear estimation using Ridge regression, *y̑*.(*t*) = *W* ∗ *x*(*t*) + *b*, where *y̑*.(*t*) is the prediction of the output, *x*(*t*) is the input, *W* and *b* are the weights and bias of the linear transformation learned from Ridge regression, respectively. All decoding was performed at 10ms bins. Decoding analyses were performed using a blocked K-fold cross validation such that all cycles from a given session were split into ten blocks, where each block was held-out from training of the decoder to generate cross-validated predictions. We used an L2 regularization constant of 10 for all analyses except for the prediction of muscle activations from smoothed spikes, where we used a value of 100 because we found higher regularization was important to achieve optimal decoding performance from smoothed spikes.

